# Trackosome: a computational toolbox to study the spatiotemporal dynamics of centrosomes, nuclear envelope and cellular membrane

**DOI:** 10.1101/2020.04.27.064204

**Authors:** Domingos Castro, Vanessa Nunes, Joana T. Lima, Jorge G. Ferreira, Paulo Aguiar

## Abstract

During the initial stages of mitosis, multiple mechanisms drive centrosome separation and positioning. How they are functionally coordinated to promote centrosome migration to opposite sides of the nucleus remains unclear. Imaging analysis software has been used to quantitatively study centrosome dynamics at this stage. However, available tracking tools are generic and not fine-tuned for the constrains and motion dynamics of centrosome pairs. Such generality limits the tracking performance and may require exhaustive optimization of parameters. Here, we present Trackosome, a freely available open-source computational tool to track the centrosomes and reconstruct the nuclear and cellular membranes, based on volumetric live-imaging data. The toolbox runs in MATLAB and provides a graphical user interface for easy and efficient access to the tracking and analysis algorithms. It outputs key metrics describing the spatiotemporal relations between centrosomes, nucleus and cellular membrane. Trackosome can also be used to measure the dynamic fluctuations of the nuclear envelope. A fine description of these fluctuations is important because they are correlated with the mechanical forces exerted on the nucleus by its adjacent cytoskeletal structures. Unlike previous algorithms based on circular/elliptical approximations of the nucleus, Trackosome measures membrane movement in a model-free condition, making it viable for irregularly shaped nuclei. Using Trackosome, we demonstrate significant correlations between the movements of the two centrosomes, and identify specific modes of oscillation of the nuclear envelope. Overall, Trackosome is a powerful tool to help unravel new elements in the spatiotemporal dynamics of subcellular structures.

## 1. Introduction

Mitosis is a highly regulated stage of the cell cycle where multiple subcellular structures take part in a complex chain of events that culminate in chromosome segregation. As cells prepare to enter mitosis, adhesion complexes disassemble (Dao et al. 2009) and the cytoskeleton reorganizes (Matthews et al. 2012; Mchedlishvili et al. 2018). At the same time, duplicated centrosomes need to migrate along the nuclear envelope so that a bipolar spindle can form (Tanenbaum and Medema 2010). This process requires the activity of multiple players, such as microtubule-associated motors kinesin-5 (Whitehead et al. 1996) and dynein (Raaijmakers et al. 2012), but also actin (Cao et al. 2010) and myosin II (Rosenblatt et al. 2004). How the dynamic changes in all these events are coordinated in space and time to ensure efficient centrosome separation and spindle assembly remains unknown.

Recent advances in live-cell imaging and image analysis techniques made it possible to access the subcellular environment and quantitatively examine its underlying mechanisms. Commercial imaging software, like Imaris and Metamorph, are equipped with automatic particle tracking functions which have been used to track centrosome pairs in 3-dimensions (Collins et al. 2014; Yamashita et al. 2015; De Simone et al. 2016). In alternative, the open-source freeware ImageJ (Eliceiri et al. 2012) with its particle tracking plugin Trackmate (Tinevez et al. 2017), has also been used for centrosome tracking in several studies (Boudreau et al. 2018; Mahen 2018; Boudreau et al. 2019). However, these generic tracking tools are not fine-tuned for the appearance and motion dynamics/constraints of centrosome pairs, which limits their tracking performance often requiring exhaustive parameter optimization, particularly when high-quality videos are not available. Moreover, when studying the dynamics of spindle formation, it is often necessary to analyze centrosomes movement in reference to the cellular and nuclear membrane (and therefore, in a non-canonical coordinates system). To the best of our knowledge, the available computational tools do not directly allow the analysis of the coordinated changes between different structures, in specific subcellular frames of reference.

The nuclear envelope (NE) also exhibits rich spatiotemporal dynamics, displaying measurable oscillations that correlate with the forces exerted by chromatin, nuclear lamina and the cytoskeleton (Chu et al. 2017; Schreiner et al. 2015; Stephens et al. 2017) and strongly influence nuclear functions (Stephens et al. 2019; Jahed and Mofrad 2019). One way to study this complex interplay between cytoskeletal forces imposed on the nucleus and the resistive forces triggered by chromatin and nuclear lamina is by measuring the dynamics of NE membrane fluctuations (Chu et al. 2017; Schreiner et al. 2015; Hampoelz et al. 2011) which allow the distinction between thermally-driven and active fluctuations (Chu et al. 2017). Importantly, during mitotic entry, chromosomes condense (Antonin and Neumann 2016) and the nuclear lamina disassembles (Georgatos et al. 1997). How these events affect the pattern of NE oscillations and impact other aspects of early spindle assembly remains to be determined. Current methods used to calculate NE fluctuations were developed under the assumption that each point of the membrane oscillates radially around its time-averaged position (Almonacid et al. 2019), (Chu et al. 2017), (Caragine et al. 2018), (Schreiner et al. 2015), (Blanchoud et al. 2010). However, describing the membrane deformations as a radial displacement is a coarse approximation that can lead to erroneous results. Also, to the best of our knowledge, there are no available toolboxes to calculate these membrane fluctuations.

Driven by these computational limitations and the need to better characterize the crosstalk between subcellular structures during mitotic entry, we developed the open-source software *Trackosome*. This novel computational tool enables a quantitative analysis of the spatiotemporal dynamics of three cellular components: centrosomes, nuclear envelope and cellular membrane. Trackosome relies on live-cell imaging datasets, where the structures of interest are independently tagged. The tool has two modules: 1) *centrosome dynamics*, used for tracking the centrosomes (or other subcellular organelles) in 3D and study their spatiotemporal relations with the nucleus and cell membrane; 2) *nuclear envelope fluctuations*, used to reconstruct, measure and analyze the dynamic fluctuations of the nuclear membrane (or other membranes) in 2D. The accurate 3D reconstruction of the centrosomes trajectories relative to the nucleus and cell membranes (in ellipsoidal coordinates) allowed us to unravel and quantify a significant correlation between centrosomes trajectories, not previously characterized. In addition, the *nuclear envelope fluctuations* module allowed us to observe and quantify distinct patterns of membrane oscillations in 2D in different stages of the cell cycle.

Trackosome is made publically available as a platform-independent MATLAB toolbox, and can be downloaded from https://github.com/Trackosome

## 2. Results

### 2.1 Dynamics of cellular reorganization during early spindle assembly

Tracking and trajectory analysis of centrosomes is performed in the Trackosome toolbox through the *centrosomes dynamics* module. This module has a graphical user interface (Figure 1A and S1) that provides useful feedback about the automatic tracking status, and allows for manual or semi-automatic corrections whenever needed. The accuracy of the tracking algorithm was tested and validated in synthetic data, created with imposed controlled conditions. The videos generated had two centrosome-like objects moving in 3D with biologically realistic dynamics (coordinates taken from real centrosomes trajectories), in a noisy environment. We set three levels of tracking difficulty by varying the signal-to-noise ratio (Figure 1B). The coordinates obtained by the Trackosome algorithm were then compared with the original coordinates used to create the videos (Figure 1C). Our tool was able to track the centrosomes with high fidelity even in highly noisy environments - signal-to-noise ratio (SNR) of 0.7 - where the particles were almost indistinguishable from the background. For all videos analyzed, the root mean squared error associated with the tracking (Figure 1C) is at the subpixel level (1 pixel = 0.189 μm).

**Figure 1.**
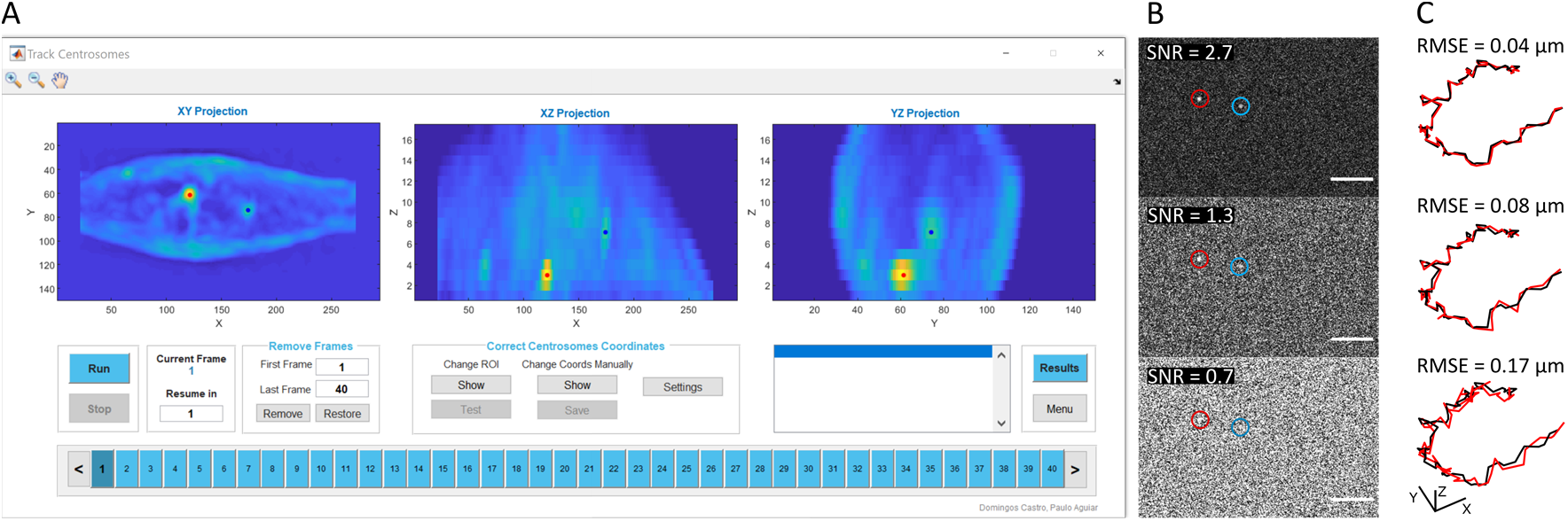
Evaluation of centrosome tracking. (A) User Interface for centrosome tracking showing the XY, XZ and YZ maximum projections for a video of a mitotic cell with the corresponding automatically identified centrosome positions (red and blue dots). (B) Frame extracted from the three synthetic videos with varying levels of SNR. Centrosomes are inside the red and blue circles. Scale bar: 10 μm. (C) Original trajectory (black) and trajectory obtained by Trackosome (red) for the centrosome on the left in B (red circle), and associated error obtained for both centrosomes. Scale bar: 1 μm.

We then recorded cells seeded on line micropatterns to normalize cell and nuclear shape (Versaevel et al. 2012). Taking advantage of the precision provided by Trackosome, we quantified specific spatiotemporal relations between three cellular structures during mitotic entry, namely centrosomes pair and nuclear and cell membranes. We chose this stage of the cell cycle as it involves extensive dynamic reorganization of the entire cell (Champion et al. 2017), providing a good benchmark to test the ability of Trackosome to detect these dynamic changes. Here, Trackosome was able to reconstruct the membranes’ surface, together with the centrosomes trajectories in 3D (Figures 2A, B). Based on their relative positions, the software was able to output different quantitative metrics, such as the distance and angles between centrosomes (Figures 2C, D), the eccentricity of the nuclear and cellular membranes (Figure 2E) and the angles between the major axis of the nucleus, cell and centrosomes (Figure 2F). These metrics describe the intracellular reorganization that occurs as cells enter mitosis, such as movement of the centrosomes to opposite sides of the nucleus (Figure 2B-D) and nucleus-centrosomes axis reorientation (Figures 2B,F), which we previously described (Nunes et al. 2020).

**Figure 2.**
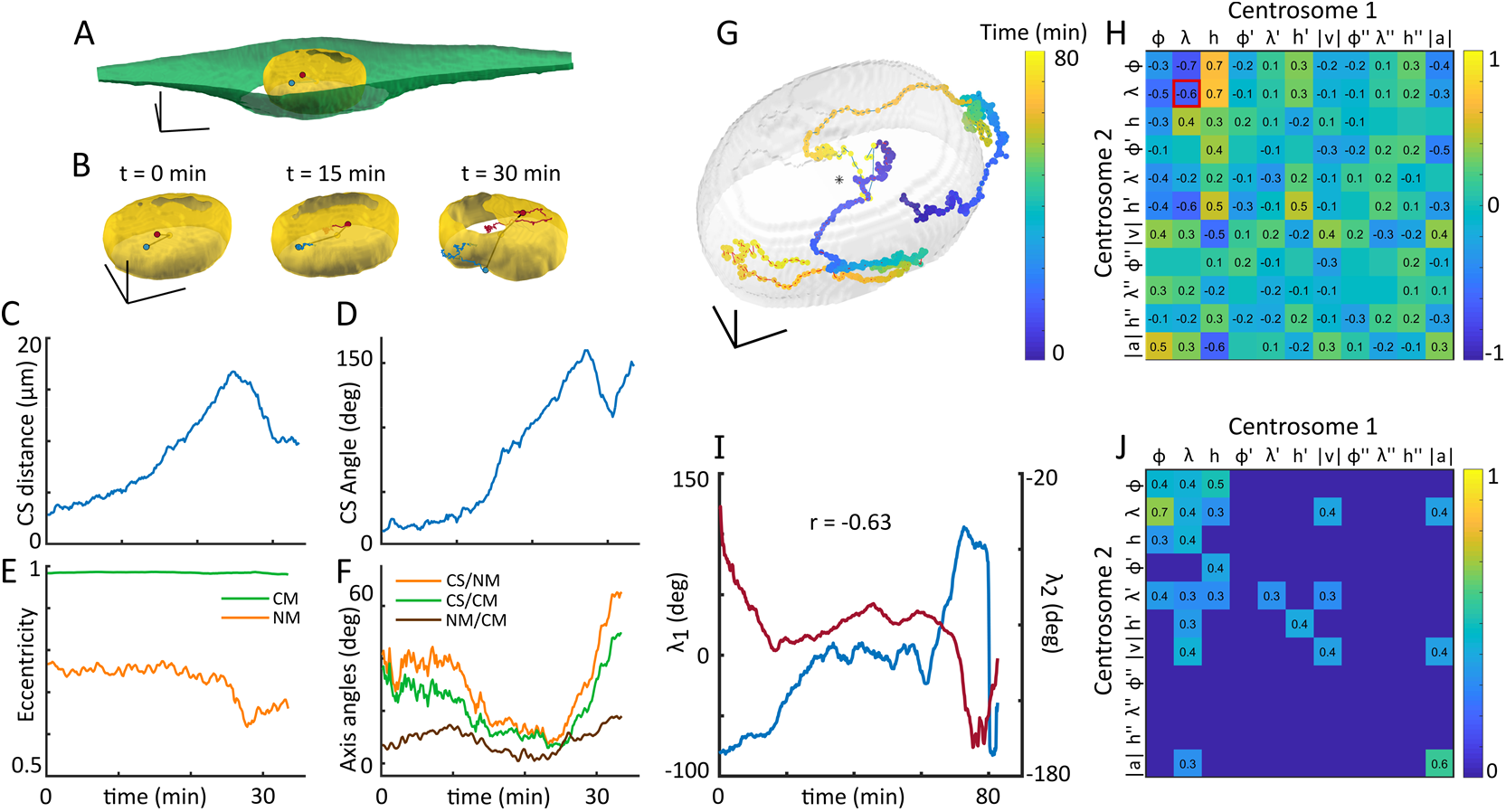
Spatiotemporal relations between cellular structures during early mitosis. (A-F) Example of Trackosome outputs for a representative cell in mitosis. (A) Three dimensional reconstruction of the cellular membrane (green), nuclear envelope (yellow) and centrosomes (red and blue dots). Scale bars: 10 μm. (B) Nuclear membrane and centrosomes at three distinct time stamps. The centrosomes trajectories (red and blue lines) evidence their migration to opposite poles of the nucleus followed by a progressive nuclear deformation. Scale bars: 10 μm. (C) Distance between centrosomes over time. The distance increases gradually during centrosome migration and decreases once the centrosomes start compressing the nucleus. (D) Angles formed between the centrosomes and the nucleus centroid over time. Note how the decrease in the distance between the centrosomes (C) occurs after centrosomes are on opposite sides of the nucleus, corresponding to the highest value for the centrosomes-nucleus angle. (E) Eccentricity of the cellular (green) and nuclear (orange) membranes evidencing that, while the cellular membrane remains morphologically stable, the nuclear membrane undergoes conformational changes after the centrosomes start deforming the nucleus. (F) Angles formed between: centrosomes axis and the nucleus major axis (orange); centrosomes axis and the cell major axis (green); the nucleus major axis and the cell major axis (brown). The centrosomes pair progressively move towards a disposition perpendicular to the major axis of the nucleus and cell. (G-J) Trajectories of centrosome pairs are spatiotemporally correlated. (G) Centrosomes positions across time (color-coded for elapsed time) surrounding the nucleus (gray). Scale bars: 5 μm. (H) Correlation matrix of the movement components (in ellipsoidal coordinates) for the trajectories shown in (G). (I) Example of the correlated features highlighted in (H, red square), corresponding to the longitude components of both centrosomes, which suggests a potential mechanical coupling between both structures. (J) Median of the absolute correlation matrices obtained from videos of different cells (n = 5), thresholded at 0.3 for visualization purposes, showing consistent correlations between centrosome trajectories across different cells.

### 2.2 Centrosomes trajectories are not independent

HeLa or U2-OS cells labelled with histone H2B-GFP/alpha-tubulin-RFP/SiR-actin or EB3-GFP/Lifeact-mCherry/SiR-DNA, respectively, were tracked until nuclear envelope breakdown (NEB). During this stage, nuclear shape remained approximately constant and correlated with the labelled chromatin, allowing us to define the median nucleus membrane (Figure 2G, grey mask) based on the 3D reconstructions of the nuclear envelope. The centrosomes, on the other hand, exhibited complex trajectories that resemble a search/adaptive path around the nucleus (Figure 2G). For this reason, to infer about the coordination of movement between the centrosomes, their trajectories were analyzed using the nucleus as a reference. An ellipsoid was fitted to the median nuclear membrane, setting the frame of reference for the new coordinate system (Figure S2). Each point of the centrosomes trajectory was defined by a latitude φ, longitude λ and height h, the respective ellipsoidal velocities φ’, λ’, h’ and velocity norm |v|, and the ellipsoidal accelerations φ’’, λ’’, h’’ and acceleration norm |a|. We calculated the correlation matrix of these features for each centrosome pair (Figure 2.H), revealing which components are temporally correlated between the two trajectories. Our results indicate a considerable degree of similarity and synchrony among trajectory pairs, with correlation values as high as 0.7, in contrast to a previous report (Waters et al. 1993). By calculating the median of the absolute correlation matrices obtained, we produced a map of the most consistent correlations (Figure 2.J). Interestingly, the positions of the centrosomes, φ, λ, h, and the acceleration norm, |a|, showed significant correlations. The latter is particularly relevant, because the acceleration of a particle reflects the force applied to it, suggesting a synchronous variation of the forces applied to both centrosomes, likely driven by kinesin-5 (Whitehead et al. 1996) or dynein (Raaijmakers et al. 2012). It is worth emphasizing that if the centrosomes movement had been described in Cartesian coordinates (i.e. ignoring the nuclear surface constraint), the observed correlations would not be evident.

### 2.3 Dynamics of nuclear envelope fluctuations driving mitotic entry

The dynamic morphology of the NE was analyzed using Trackosome’s *membrane fluctuations* module (Figure S4). The membrane oscillations are determined by calculating the orthogonal displacement of each point of the membrane with respect to its medial position (Figure S5). Our method does not rely on prior assumptions regarding the nucleus shape. We believe that this approach leads to a more realistic description of the membrane displacements than the radial displacement approximations (with respect to the membrane’s centroid) usually done in the literature (Chu et al. 2017; Caragine et al. 2018; Schreiner et al. 2015; Blanchoud et al. 2010; Almonacid et al. 2019). The radial displacements are particularly flawed for irregularly shaped nucleus with wide fluctuations (Figure 3A), which generally limits its use to cells in interphase. Here we were able to quantify and compare the nuclear deformations for cells in interphase and mitosis.

As before, we recorded cells seeded on line micropatterns. From our analysis, it is possible to confirm that in interphase, cells present subtle but measurable nuclear membrane movements (Figure 3B, C). As a negative control, fluctuation measurements were made for interphase cells fixed with formaldehyde, which showed a significant decrease of undulations upon fixation (Figure 3.B, C).

**Figure 3.**
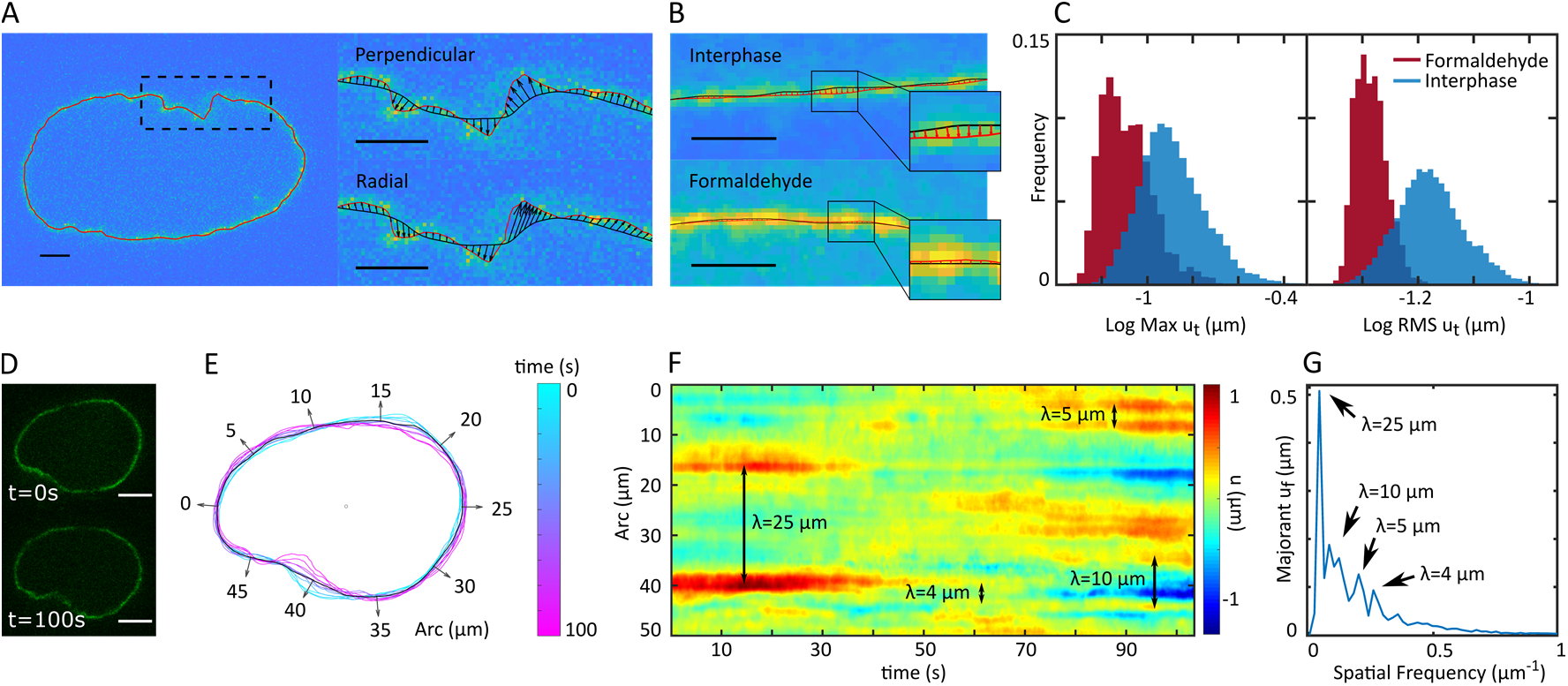
Nuclear membrane fluctuations captured with Trackosome. (A-C) The perpendicular membrane displacements measured with Trackosome are sensible to subtle membrane movements. (A) Membrane segmentation (red) of a representative nucleus in prophase (left) and a detailed view of the upper region of the membrane (right) illustrating the difference between defining the fluctuations (black vectors) as perpendicular (top right) or radial (bottom right) movements of the current membrane (red) around the median membrane (black). For the radial displacements, the centroid of the median membrane is used as origin. Perpendicular displacements offer a more realistic description of the membrane movement around its basal position. Scale bars: 2 μm. (B) Representative examples of the nuclear membrane fluctuations (red vectors) of a cell in interphase (top) and a cell fixed in formaldehyde (bottom). The small fluctuations measured for the cell in interphase are clearly larger than those obtained for the fixed cell, suggesting that they are the result of subtle membrane movements. (C) Distributions of the maximum fluctuation per frame, Max *u_t_* (left), and the root mean square of the fluctuation per frame, RMS *u_t_* (right), for cells in interphase (44 cells with approximately 600 frames each) and formaldehyde (4 cells with approximately 500 frames each). With logarithmic scales, the distributions are approximately Gaussian and easily distinguishable. (D-G) Typical nuclear fluctuations results obtained with Trackosome for a cell in prophase. (D) Two frames of the original video. Scale bar: 5 μm. (E) Centered nucleus membrane extracted from the video in (D) at different time stamps (colored scale), with vectors (gray) indicating the arc in micrometers around the median membrane (black), pointing in the direction along which the fluctuations are calculated. (F) Map of the fluctuations amplitude, *u*, obtained for all the frames. The y axis corresponds to the arc around the median membrane, marked by the vectors in (E). (G) Majorant of frequency dependent fluctuations, *u_f_*, obtained by finding the maximum amplitude of the spatial Fourier Transform (FT) for each frequency across frames. This FT curve shows the maximum fluctuation amplitude of each wavelength, thus, its peaks reveal the amplitude and wavelength of the most significant curvatures of the nuclear envelope, evidenced both in (F) and (G).

During mitotic entry, chromosomes condense (Antonin and Neumann 2016) and the nuclear lamina disassembles (Georgatos et al. 1997). Whether these events change nuclear behavior remains to be determined. We analyzed nuclear membrane fluctuations of cells using Trackosome (Figure 3D-G). After segmenting and registering (centering on centroid) the NE (Figure 3E), the fluctuations *u* of each frame are calculated assuming normal displacements (Figure 3A, top). The values obtained were concatenated frame-by-frame and filtered in time and space with a 2D Gaussian kernel. The resulting fluctuations map reveals how the envelope curvatures are dominated by specific wavelengths (Figure 3F). These wavelengths are determined by calculating the spatial Fourier Transform (FT) of the fluctuations, *u_f_*, for each frame and then obtaining the maximum magnitude of each wavelength across frames (Figure 3G). Calculating the majorant for each frequency component *u_f_* has the advantage of highlighting the most well defined spatial frequencies, even if they occur for a limited number of frames. On the other hand, averaging the FT curves of all frames would attenuate these components and the magnitude of the final FT peaks would not correspond to the magnitude of the actual membrane waves that originated them, thus providing a less intuitive readout.

The FT curve allowed us to identify and quantify the most prominent spatial frequencies and obtain relevant information about the dynamics of NE fluctuation. For the cell in prophase exemplified in Figure 3D-G, the highest FT peak, with a wavelength of 25 μm (half of the nucleus perimeter), is the result of the large membrane displacement seen around the 15 and 40 μm arc landmarks in Figure 3E, F. As prophase progresses, exertion of compression forces in these two separate points makes the nuclear membrane wobble with a wavelength which is half of the membrane perimeter (Figure S5). In addition, the FT analysis allowed the identification of membrane “wrinkles” with a spatial signatures defined by wavelengths of approximately 10, 5 and 4 μm (Figure 3F,G). Trackosome captures these spatial signatures with high accuracy even for curvatures with amplitudes at the subpixel level under noisy backgrounds (Figure S5).

To help uncover the processes behind these observed spatial frequencies, membrane fluctuations were calculated for cells in early mitosis and interphase (Figure 4A,B). We calculated the majorant *u_f_* for all cells (as in Figure 3G) and then the median curve across trials for each group. Our results show a significant difference in the FT curves of the two groups (Figure 4B). In interphase, the magnitude of the fluctuations is very low through the entire spectrum of wavelengths, indicating that the membrane barely deviates from its basal position. Still, the *u_f_* magnitude is above the noise floor stablished by the formaldehyde curve. This behavior changes in prophase, as there is an increase of the fluctuations amplitude for the entire range of wavelengths. This increase is particularly significant for the low frequency components, reflecting the occurrence of nucleus-wide deformations as described above.

**Figure 4.**
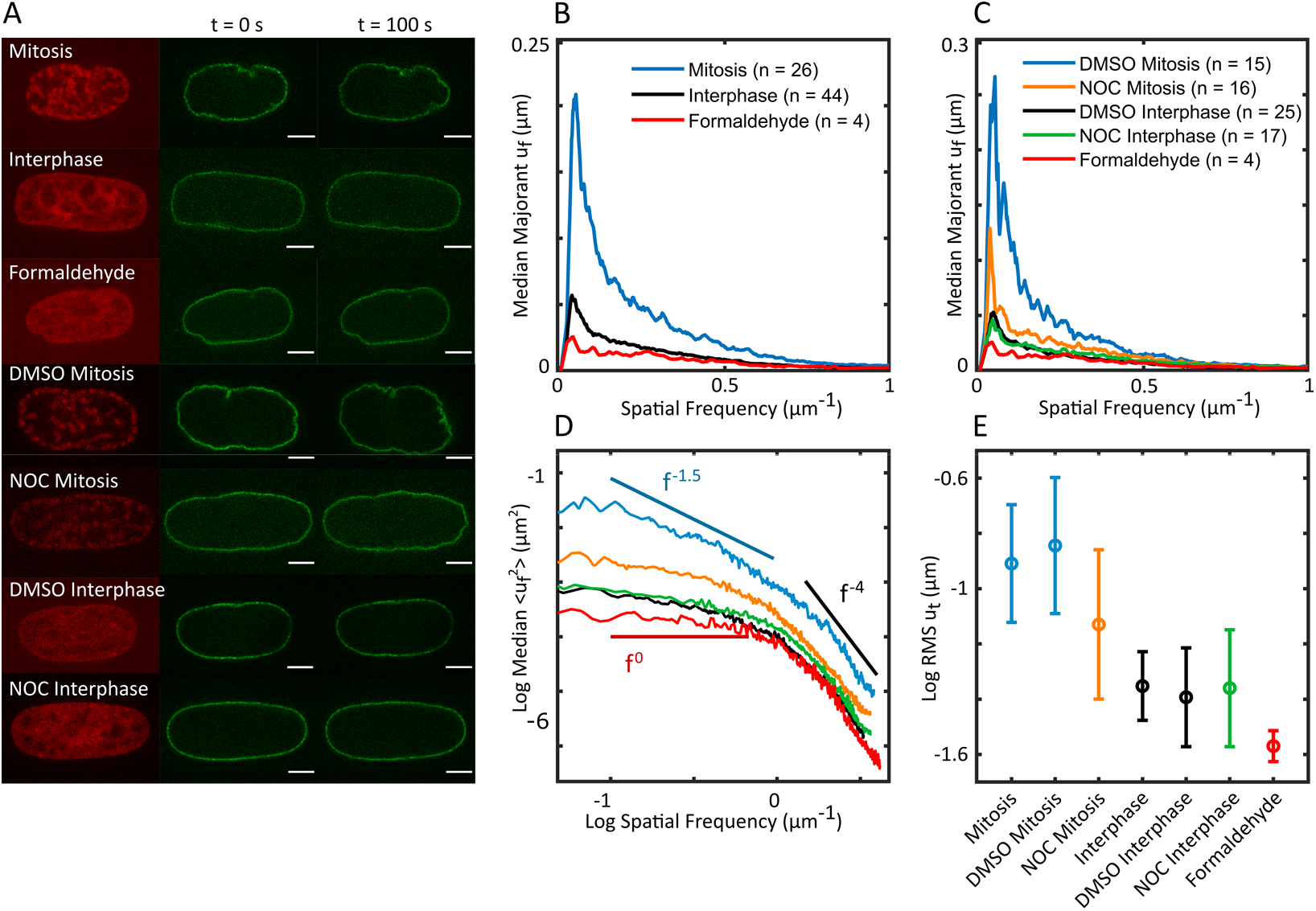
Nuclear membrane fluctuations vary with the stage of the cell cycle and the physiological treatment. (A) Representative nucleus of each group. The phase of the cell cycle is evidenced by the marked histone (red), taken from the first frame of each video. The nuclear envelope (green) is shown at two different times stamps to illustrate the degree of membrane undulations in each group. Scale bar: 5 μm. (B) Median of the majorant frequency dependent fluctuations, *u_f_*, obtained for groups of cells in interphase and early mitosis. The curve for cells fixed with formaldehyde was also included to set the noise limit. (C) Median of the majorant *u_f_* obtained for groups of cells in interphase and mitosis, treated with DMSO, nocodazole (NOC) and fixed with formaldehyde. NOC caused a significant decrease of the membrane fluctuations in mitosis. (D) Median across cells of the average FT of the squared fluctuations of each cell, <*u_f_^2^*>, for the groups represented in (C). In logarithmic scales, the <*u_f_^2^*> curves show regions dominated by different frequency dependencies, limited by the solid lines with slopes f^0^, f^−1.5^ and f^−4^. (E) Mean and standard deviation of the log RMS of the fluctuations for all the tested groups. Using a logarithmic scale, the RMS fluctuations distributions are approximately normal and can thus be described by their mean value and standard deviation.

Fluctuations of the NE are known to depend on the cytoskeleton and, in particular, on the microtubule (MT) network (Chu et al. 2017; Hampoelz et al. 2011). To assess whether Trackosome was sensitive to detect these changes, we treated cells with low doses of nocodazole (NOC) to suppress the polymerization of MTs without affecting overall cell structure. Interphase cells treated with NOC had an FT curve similar to the control curve obtained for interphase cells treated with DMSO only (Figures 4A, C). On the other hand, nuclear fluctuations of mitotic cells were significantly altered with the addition of NOC. The FT obtained for this experimental condition showed a decrease in the low-frequency peak when compared with the control group. This suggests that the disruption of the MT cytoskeleton led to a decreased nuclear compression, which is in agreement with the fact that the MTs induce large-scale deformations of the nucleus envelope (Hampoelz et al. 2011; Schreiner et al. 2015). The frequency components beyond the peak were also severely attenuated, which means that the shorter membrane “wrinkles” were mitigated as well. Most of these effects can be qualitatively evaluated examining the nuclei images directly (Figure 4A), but Trackosome provides a precise quantification.

Biological membranes exhibit passive fluctuations, thermally excited at physiological temperatures. The equilibrium properties of these undulating systems can be modeled by Helfrich-type models (Helfrich 1978). To evaluate the spectral dependencies in line with Helfrich theory, we calculated the average FT of the squared fluctuations for each cell, <*u_f_^2^*>, and then the median curve across cells for each group. The curves of the fluctuations obtained for the NOC and control groups show regions with distinct spectral dependencies (Figure 4D). For frequencies above 1.5 μm^−1^ (wavelengths below 0.7 μm) the FT curves follow a f^−4^ power law, consistent with what is expected for thermally driven fluctuations at short wavelengths (Brandt et al. 2011). At lower frequencies, each group follows a different power law, ranging from close to f^0^ for formaldehyde, up to f^−1.5^ for the control group in mitosis. These can be attributed to different nonequilibrium active forces governing each system. It is important to note that in Helfrich theory these spectral dependencies are defined in terms of the adimensional wavenumber *q*, and not spatial frequency. However, for the frequency ranges marked by the solid lines in Figure 4D, the slopes are equivalent in both scales (Figure S7).

Finally, we compared the fluctuations amplitude (in terms of Log RMS *u_t_* as in Figure 3C right) for all the groups (Figure 4E). NOC led to a significant decrease on the fluctuations amplitude during prophase. Importantly, this decrease was not caused by the vehicle DMSO, since prophase cells in the presence of DMSO had higher fluctuations than the untreated prophase cells. Also, the fluctuations obtained for the interphase cells are similar among the different groups and the range of amplitudes is consistent with what was already described in the literature (Schreiner et al. 2015).

Overall, our results indicate that nuclear membrane fluctuations increase as cells transition from interphase to mitosis and that interfering with the MT network significantly reduces the large scale deformations of the nucleus during this stage.

## 3. Discussion

Recent developments in light microscopy have generated extensive datasets that allow unprecedented access to subcellular events with high spatiotemporal resolution. Here, we report on Trackosome, a new open-source software that enables an automated quantitative analysis of the spatiotemporal dynamics of subcellular structures even in conditions of low SNR. During the transition from G2 to mitosis, chromosomes condense (Antonin and Neumann 2016), cells round up (Matthews et al. 2012) and centrosomes separate (Whitehead et al. 1996). Under this context, the extent of centrosome separation at the time of NEB remains a matter of debate. While some reports show that cells can enter mitosis with unseparated centrosomes (Kaseda et al. 2012; Whitehead et al. 1996), others indicate that centrosomes are fully separated at NEB (Magidson et al. 2011; Mardin et al. 2013; Nunes et al. 2020). These discrepancies highlight the need to carefully determine how the events leading up to mitosis are coordinated to ensure efficient spindle assembly. By using the different modules available on Trackosome, we were able to accurately correlate centrosome movement and centrosome-nucleus axis reorientation during mitotic entry. Our results, using cells seeded on fibronectin micropatterns, indicate that centrosomes are fully separated before NEB and that their movement is coordinated to optimize their positioning on opposite sides of the nucleus (Nunes et al. 2020), contrarily to previous observations (Waters et al. 1993). This coordination was evidenced using elliptical coordinates to describe the centrosome trajectories around the nucleus. Interestingly, the high correlation scores between movement components of both centrosomes strongly indicates a mechanical coupling between the two structures. This coupling could be provided by specific pools of dynein on the cell cortex (Kotak et al. 2012; Woodard et al. 2010) or at NE (Bolhy et al. 2011; Splinter et al. 2012), which are known to generate pulling forces on centrosomal microtubules to position asters (Laan et al. 2012). Determining the exact nature of this mechanical coupling between the centrosomes during mitotic entry will be of interest in the future.

The mechanical properties of the nucleus depend on the chromatin condensation state (Stephens et al. 2017; 2018) and the nuclear lamina (Lammerding et al. 2006). Given that Lamin A levels (Chu et al., 2017; Moir et al. 2000) and chromatin compaction (Hinde et al. 2012) change throughout the cell cycle, it is possible that the mechanical properties of the nucleus change accordingly. In agreement, nuclear envelope fluctuations, which reflect forces imposed on the nucleus, were shown to vary depending on the cell cycle stage (Chu et al. 2017). However, whether the mechanical properties of the nucleus change during the transition from G2 to mitosis, when mitotic chromosomes are condensing and the nuclear lamina disassembles, remains unclear. Here, we report on a new tool to analyze NE fluctuations during mitotic entry that measures orthogonal displacements relative to the medial position of the nuclear membrane. We reveal that chromosome condensation, together with microtubules, trigger significant changes in the spatial pattern of NE fluctuations as cells prepare to enter mitosis. To the best of our knowledge, it is the first time that the transition from G2 to prophase is characterized in terms of nuclear envelope fluctuations. In the future, it will be interesting to determine how these fluctuations reflect the mechanical properties of the nucleus.

The algorithms compiled in the computational toolbox Trackosome provide a reliable and accurate instrument to help uncover new elements in the spatiotemporal dynamics of subcellular structures. Importantly, this toolbox has the potential to be adapted to other experimental conditions such as the study of cell migration and cell polarity, where the capacity to analyze dynamic datasets with high accuracy is highly relevant.

## Acknowledgments

The authors would like to thank Margarida Dantas for the critical reading of the manuscript and Helder Maiato for providing access to microscopes and equipment. The authors would like to thank Katharine Ullman for the HeLa POM121-3x-GFP/H2B-mCherry. This work was funded by grants from FEDER - Fundo Europeu de Desenvolvimento Regional funds through the COMPETE 2020 - Operacional Programme for Competitiveness and Internationalization (POCI), Portugal 2020, and by Portuguese funds through FCT - Fundação para a Ciência e a Tecnologia/Ministério da Ciência, Tecnologia e Ensino Superior in the framework of the project PTDC/BEX-BCM/1758/2014 (POCI-01–0145-FEDER-016589). V.N. is supported by grant PD/BD/135545/2018 from the BiotechHealth FCT-funded PhD program. J.T.L. is supported by grant SFRH/BD/147169/2019 from FCT.

## Author contributions

D.C. and P.A. developed all MATLAB computational tools. V.N., J.T.L. and J.G.F. performed experiments. J.G.F. and V.N. designed the experiments. The manuscript was written primarily by D.C., P.A. and J.G.F‥ J.G.F. obtained funding and provided resources.

## 4. Materials and Methods

### 4.1 Biological methods

#### Cell culture

Cell lines were cultured in Dulbecco’s Modified Eagle Medium (DMEM; Life Technologies) supplemented with 10% fetal bovine serum (FBS; Life Technologies) and grown in a 37°C humidified incubator with 5% CO2. HeLa POM121-3xGFP/H2B-mCherry cell line was a kind gift from Katharine Ullman. HeLa cell line expressing histone H2B-GFP/mRFP-α-tubulin was generated in our lab using lentiviral vectors.

#### Micro-patterning

Micro-patterns to control individual cell shape and adhesion pattern were produced as previously described (Azioune et al. 2009). Briefly, glass coverslips (22 × 22mm No. 1.5, VWR) were activated with plasma (Zepto Plasma System, Diener Electronic) for 1 min and incubated with 0.1 mg/ml of PLL(20)-g[3,5]-PEG(2) (SuSoS) in 10 mM HEPES at pH 7.4, for 1 h, at RT. After rinsing and air-drying, the coverslips were placed on a synthetic quartz photomask (Delta Mask), previously activated with deep-UV light (PSD-UV, Novascan Technologies) for 5 min. 3 μl of MiliQ water were used to seal each coverslip to the mask. The coverslips were then irradiated through the photomask with the UV lamp for 5 min. Afterwards, coverslips were incubated with 25 μg/ml of fibronectin (Sigma-Aldrich) and 5 μg/ml of Alexa546 or 647-conjugated fibrinogen (Thermo Fisher Scientific) in 100 mM NaHCO3 at pH 8.6, for 1 h, at RT. Cells were seeded at a density of 50.000 cells/coverslip and allowed to spread for ~10-15h before imaging. Non-attached cells were removed by changing the medium ~2h-5h after seeding.

#### Time-lapse microscopy

For time-lapse microscopy, 12-24 h before the experiments, 5×10^4^ cells were seeded on coverslips coated with FBN (25μg/ml; F1141, Sigma). Prior to each experiment, cell culture medium was changed from DMEM with 10% FBS to Leibovitz’s-L15 medium (Life Technologies) supplemented with 10% FBS and Antibiotic-Antimycotic 100X (AAS; Life Technologies). When SiR-dyes were used, they were added to the culture medium 30min-1h before acquisition (20nM Sir-tubulin or 10nM Sir-DNA; Spirochrome). Where stated, nocodazole was added to the cells 30 min before the experiment (20 nM; Sigma-Aldrich). Live-cell imaging was performed using temperature-controlled Nikon TE2000 microscopes equipped with a modified Yokogawa CSU X1 spinning-disc head (Yokogawa Electric), an electron multiplying iXon+ DU-897 EM-CCD camera (Andor) and a filter-wheel. Three laser lines were used for excitation at 488, 561 and 647nm. For nuclear membrane fluctuations, an oil-immersion 100x 1.4 NA Plan-Apo DIC (Nikon) was used. All the remaining experiments were done with an oil-immersion 60x 1.4 NA Plan-Apo DIC (Nikon). Image acquisition was controlled by NIS Elements AR software. For centrosome tracking, 17-21 z-stacks with a 0.5μm separation were collected every 20 sec or, to analyze centrosome correlations, every 10 sec. For nuclear envelope fluctuation measurements, a single z-stack was collected every 100 msec.

### 4.2 Trackosome toolbox: Centrosome Dynamics

#### Tracking algorithm

A 3D Laplacian of a Gaussian filter is applied to the centrosomes video to highlight centrosome-like blobs. The user must then select the approximate position of the two centrosomes in a XY maximum projection of the first frame. These are the seeding points for the tracking loop that follows. For each frame, two 3D regions of interest (ROI) are centered at the coordinates of the centrosomes obtained for the previous frame (for the first frame, it uses the coordinates selected by the user). The ROI are large enough to accommodate the centrosomes dislocations in-between frames. Each ROI is thresholded to isolate the regions of high intensity, and then iteratively shortened until it confines a peak of intensity with the dimensions of the centrosome. Once each centrosome is enclosed by a mask, the centroids are found by projecting the masked intensities along the three axis and fitting a Gaussian to each intensity profile – the coordinates of the centroid correspond to the peaks of the fitted Gaussians (Figure S3).

#### Nuclear and cellular membrane reconstruction

The images from each video (nucleus and cellular membrane) are binarized and submitted to an iterative set of morphological operations intended to create a closed, smooth, binary volume. The membrane of each structure corresponds to the outer pixels of their binary volume. The major axis of the cell and nucleus are obtained by calculating the principal components of the binary volumes for each frame.

### 4.3 Correlation of centrosomes trajectories

The correlations among centrosomes displacements were evidenced by defining their movement as two ellipsoidal trajectories surrounding the approximately ellipsoidal nucleus (Figure S3). The nucleus was centered assuring a common centroid among frames. The median nucleus was obtained by calculating the median binary volume across time. The videos were analyzed until the nuclear envelope breakdown. An ellipsoid was fitted to each median nucleus, setting the coordinate system used to define the ellipsoidal trajectories. The trajectories of the centrosomes were normalized with regards to the centroid of the nucleus for each frame. The transformed trajectories were converted to ellipsoidal coordinates, using the fitted ellipsoid as a referential of the coordinate system. The trajectories were also characterized by the norm of the ellipsoidal velocity and acceleration, and their decomposition along the latitude, longitude and altitude directions at each point. Once all the features of both centrosomes trajectories are calculated, the correlation matrix can be directly obtained. We only included videos with long and changing centrosome trajectories, since short monotonic trajectories would invariantly have high, but misleading, correlation coefficients. We also excluded videos with non-stationary nucleus because a moving referential would lead to fallacious correlation values.

### 4.4 Trackosome toolbox: membrane fluctuations

We consider that the points of the membrane move in a direction perpendicular to its surface. Therefore, we define the fluctuations as being the distance from each point of the median membrane to the membrane at a given frame, along a direction normal to the median membrane. The membranes are segmented for all frames and then centered assuring a common centroid among frames. To calculate the basal position (the reference membrane) of the nuclear envelope, we calculated the median projection of the centered frames and segmented the resulting membrane. Each point of the reference membrane is associated with a normal vector which defines the direction of the membrane displacements. The fluctuations are then measured by calculating the distance from the reference membrane to the membrane of each frame, along the directions defined by the normal vectors (Figure S5).

## Supplementary Information

**Figure S1.**
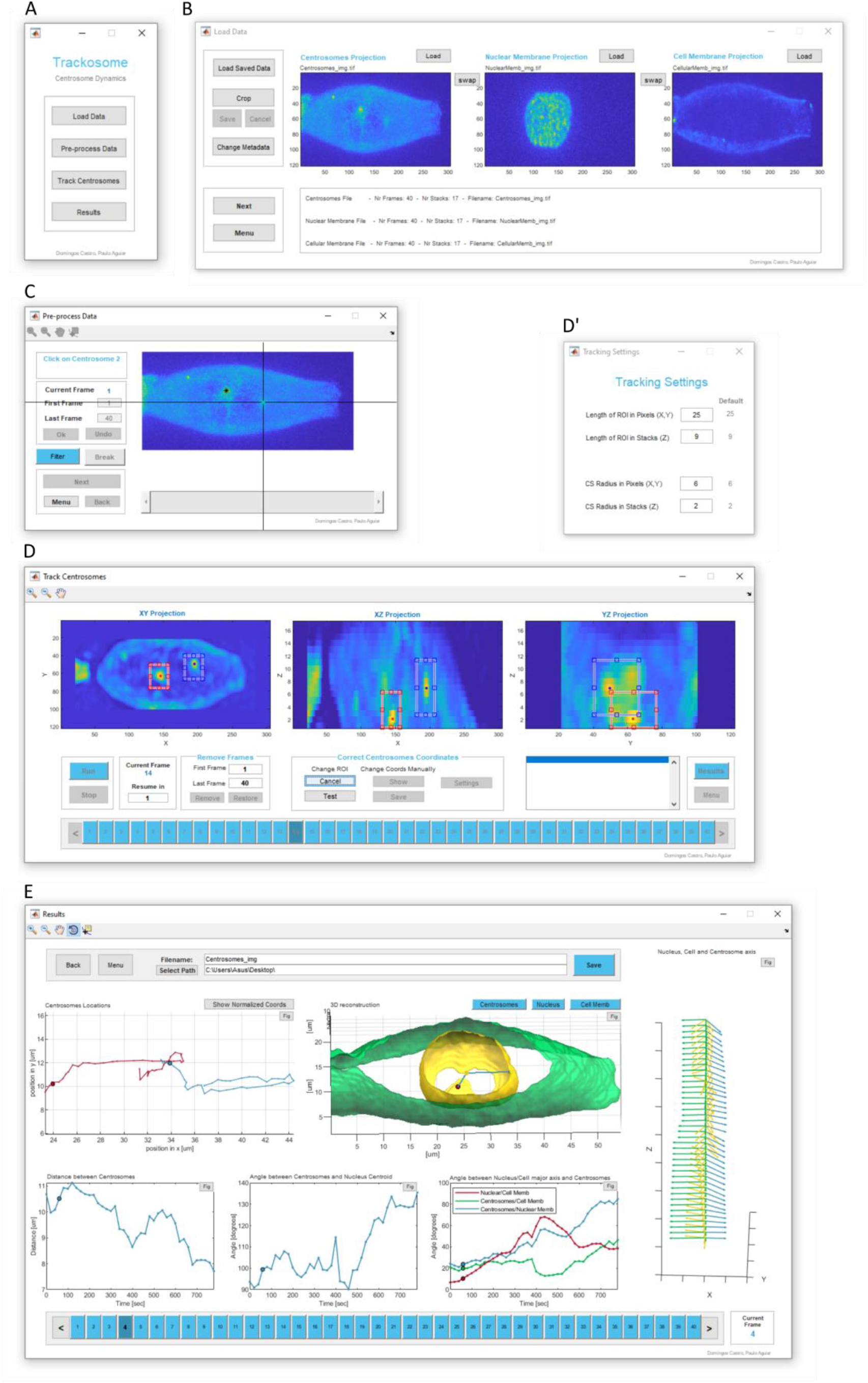
Trackosome User Interface: Centrosome Dynamics module. (A) Main menu to open the windows shown in B, C, D and E. (B) Load data window, where the user can import the 3D live-cell imaging videos from the centrosomes, nuclear membrane and cellular membrane (.tiff, .mat, .nd2). It is also possible to load .mat files previously exported by Trackosome. These populate the entire interface with the data contained in the file, allowing the user to reexamine previous analysis. (C) Pre-processing window to filter the centrosomes channel, trim the videos, and select the approximate initial coordinates of the centrosomes. (D) Centrosomes tracking window. The user can follow the tracking results for each frame in real-time. The algorithm can be stopped at any time to correct eventual mistakes. There are two modes of coordinates correction: 1) “Change ROI”: move and resize the regions-of-interest to guarantee that they contain the centrosomes (option currently selected in the shown image); 2) “Change Coords Manually”: manual selection of the new 3D coordinates of a centrosome. The algorithm can proceed from the corrected frame. These corrections can also be done after analyzing the full video. The array of blue buttons at the bottom allow the user to navigate between frames and provide visual feedback regarding the tracking status of each frame: blue - ok; yellow - problem finding the centrosomes; red - forced break due to error; gray - coordinates manually changed. The user can also discard specific frames from the analysis. (D’) Window to change the main settings of the tracking algorithm. (E) Results window where the user can inspect and save the results obtained. The results can be exported as .xlsx, .csv files and also a .mat file that stores all the data from the interface. The .mat file allows the user to reload the full Trackosome interface with the stored data. Also, the user can access relevant stored variables (such as the membrane reconstructions) by loading this .mat file directly on the Matlab command window.

**Figure S2.**
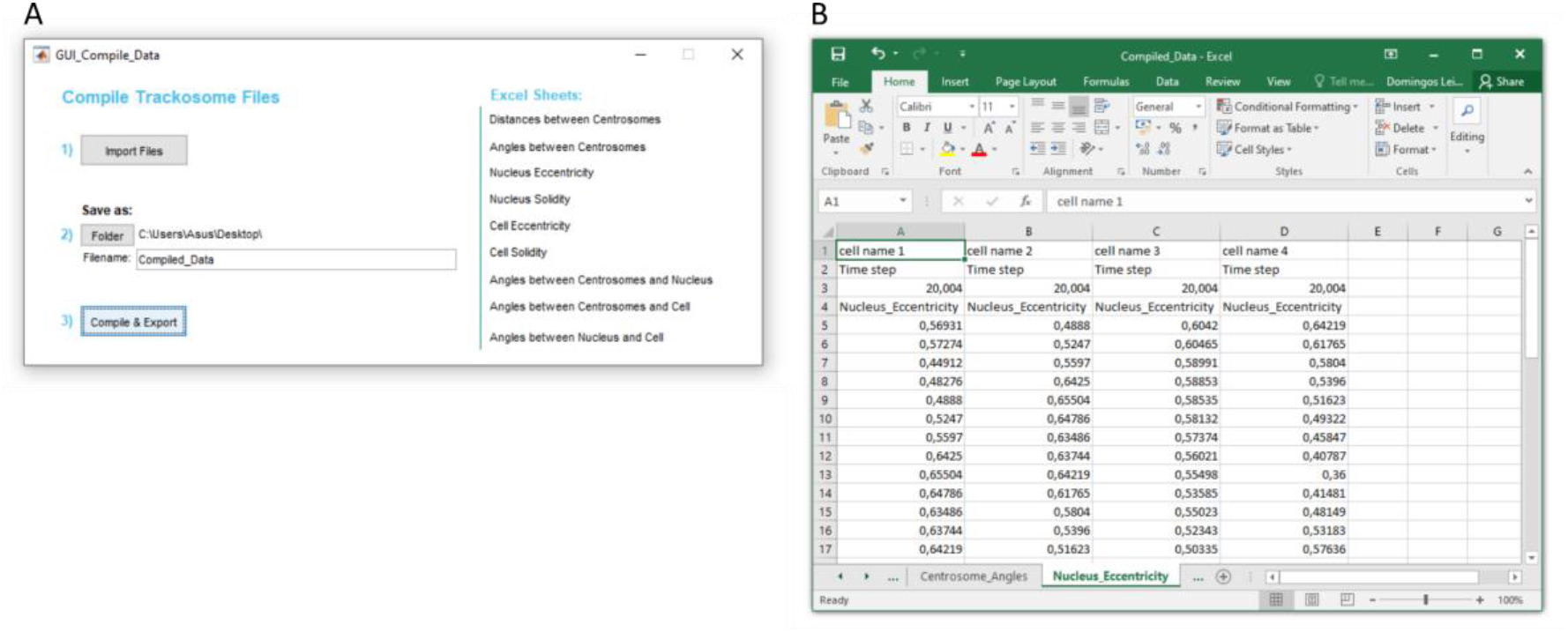
Compile Data module. (A) The toolbox includes a separate module to compile data from different .csv files exported by Trackosome. (B) Resulting excel sheet with the data compiled from different videos. Each metric is attributed to a different sheet. The data of each file (filename, time step in seconds and metric values) are concatenated side-by-side in each metric sheet.

**Figure S3.**
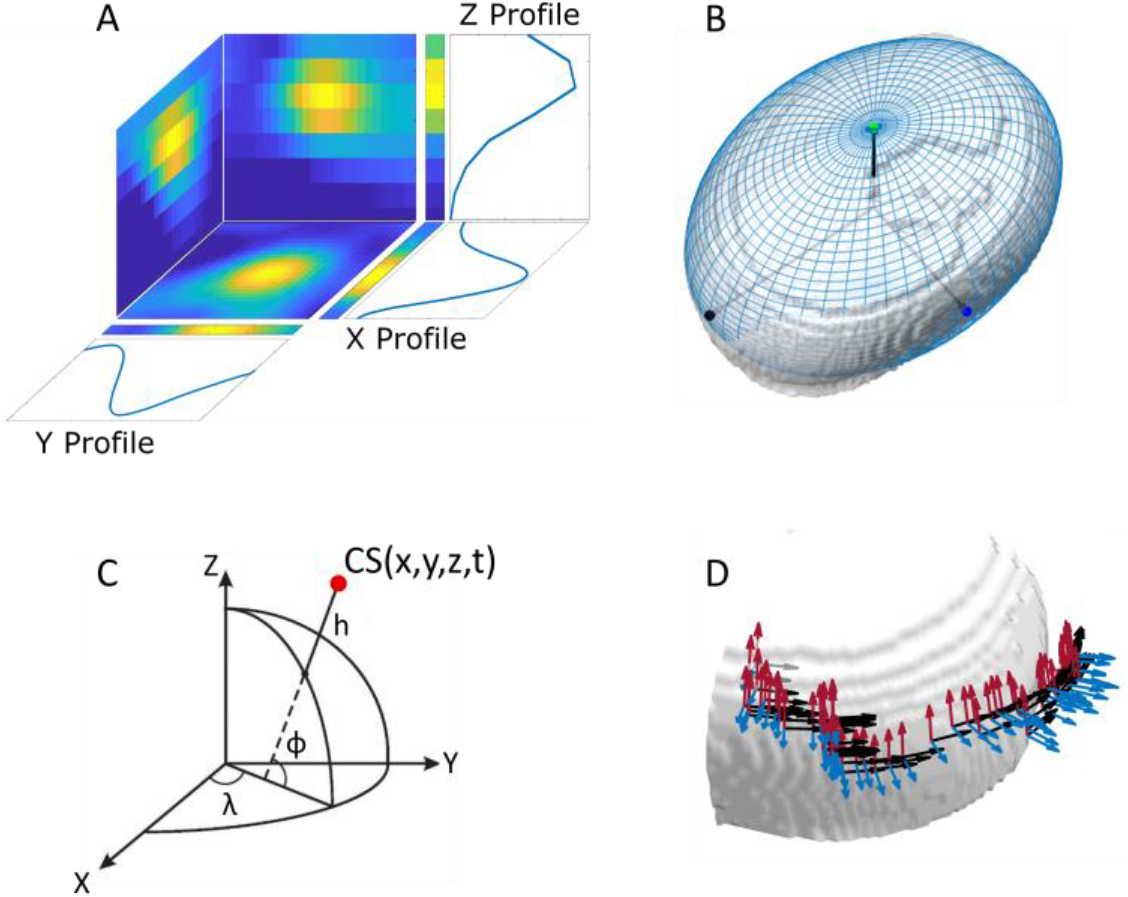
Centrosome tracking with Trackosome and definition of elliptical coordinates to unravel trajectory correlations. (A) Intensity projections of the initial region-of-interest (ROI) obtained for a centrosome at a given frame. The ROIs are centered in the coordinates of the centrosome obtained in the previous frame. Each dimension of the initial ROI is iteratively shortened until it confines a blob with the dimensions of the centrosome. The final 3D coordinates of the centrosome are found by fitting a Gaussian curve to the intensity profiles along the X, Y and Z axis. The (x,y,z) position of the centrosome corresponds to the mean value of each Gaussian. (B) Ellipsoid (blue) fitted to the median nucleus (grey) used as referential for the ellipsoidal coordinates. (C) Conversion from cartesian (x,y,z) to ellipsoidal (ϕ, λ, h) coordinates. (D) Each point of the centrosomes trajectories is associated with an orthonormal basis defined by a latitude versor (red), longitude versor (black) and height versor (blue). These indicate the direction along which the ellipsoidal velocities and accelerations are calculated for each position.

**Figure S4.**
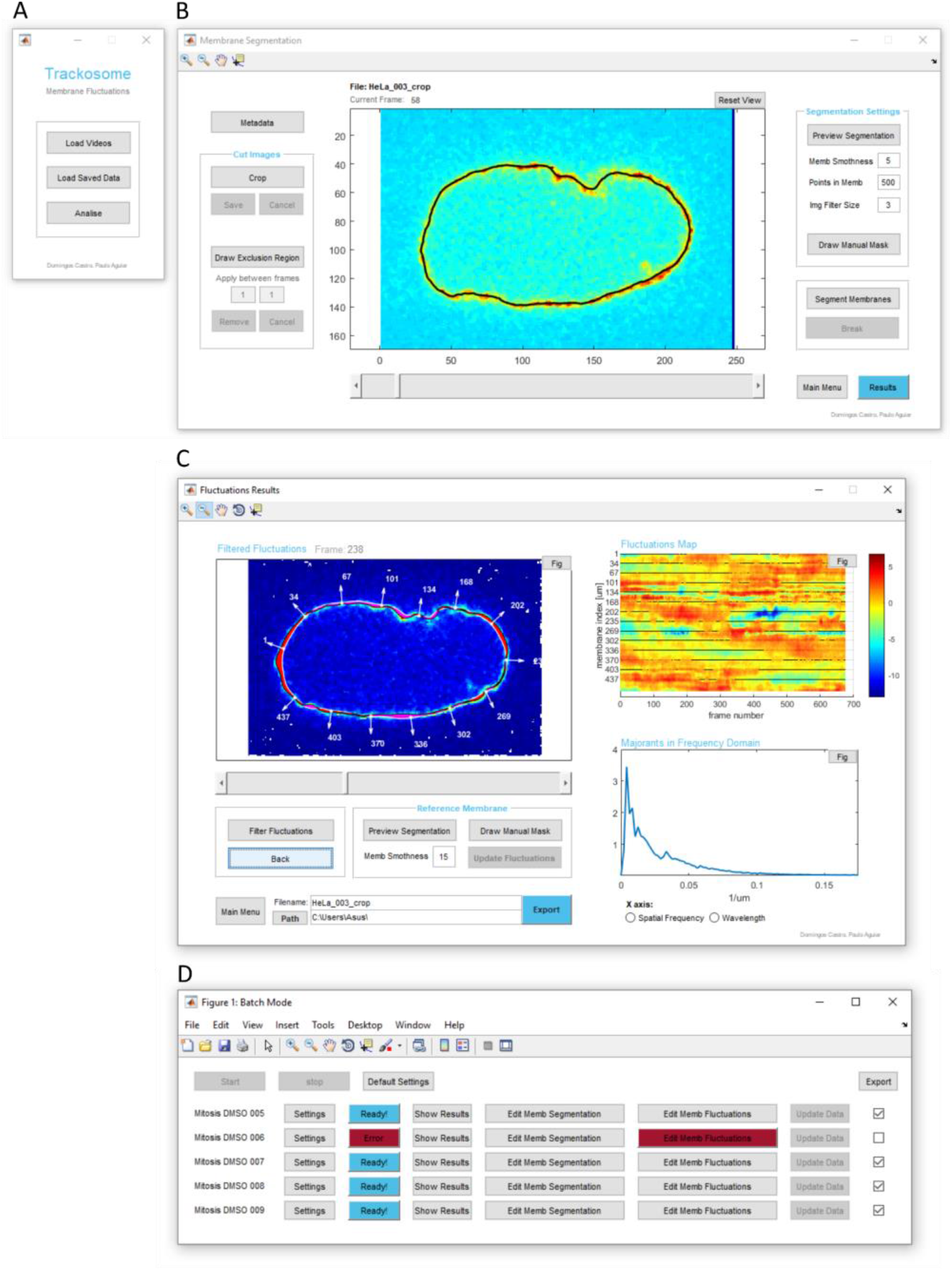
Trackosome User Interface: Membrane Fluctuations module. (A) Main menu to load the 2D videos (.tiff, .mat, .nd2). The loading function used is *uipickfiles* (https://www.mathworks.com/matlabcentral/fileexchange/10867-uipickfiles-uigetfile-on-steroids). If only one video is selected, the user is directed to the window shown in B. If multiple files are selected, the user is directed to batch analysis mode, shown in D. It is also possible to open previously exported files and load the entire interface with the imported data. (B) Window to perform membrane segmentation. The user can edit the video before segmenting the membranes. The edit options include cropping the frames, drawing masks to guide membrane segmentation, and removing manually drawn regions from specified frames (to eliminate, for example, high intensity noise blobs located near the membrane). It is possible to preview the segmentation of any frame to optimize parameters before starting the segmentation of the entire video. (C) Results window where the user can inspect, correct and export the results obtained. The corrections include editing the reference membrane and adjusting the spatiotemporal filters applied to the membrane fluctuations. (D) Window for batch analysis, opened if the user selects multiple videos in the Main Menu. In the window shown, five files were selected. Each file is associated with a set of buttons to edit parameters, show the results obtained, edit the membrane segmentation (opens window B), edit fluctuation results (opens window C), and activate/deactivate exporting.

**Figure S5.**
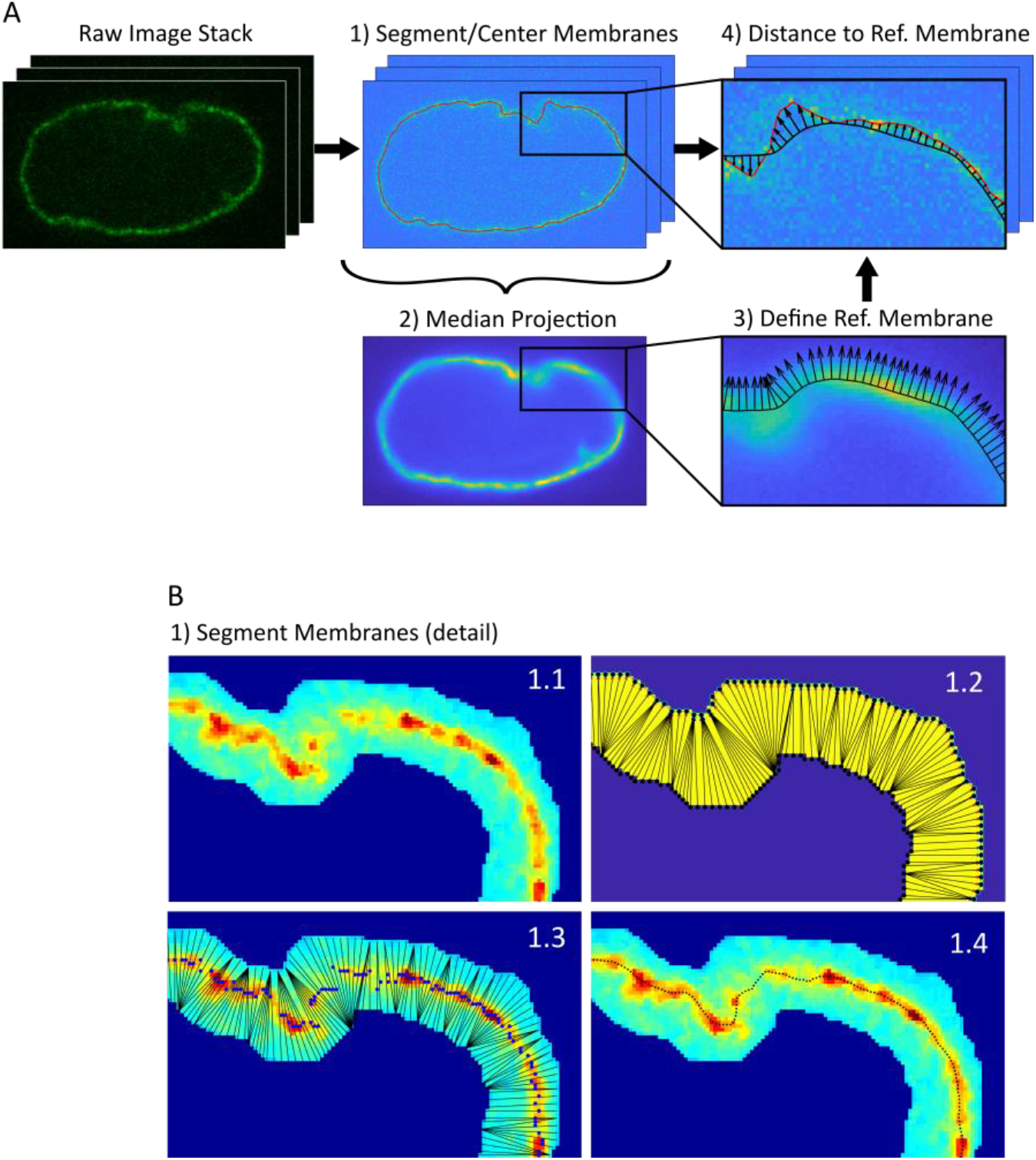
Nuclear membrane fluctuations algorithm. (A) Overview of the four main steps of the algorithm: 1) membrane segmentation and centering; 2) median projection of the centered frames; 3) segmentation of the membrane obtained in 2, defining the reference membrane and the associated normal vectors; 4) calculating the membrane fluctuation for each frame as the distance between the reference membrane and the membrane of the current frame, along the direction defined by the vectors orthogonal to the reference membrane. (B) Detail of the membrane segmentation algorithm for a given frame: 1.1) filter and mask the image; 2) the points that constitute the two borders of the mask are connected by the shortest segment; 3) find the pixel with maximum intensity for each segment defined in 1.2; 4) filter the positions of the pixels found in 1.3.

**Figure S6.**
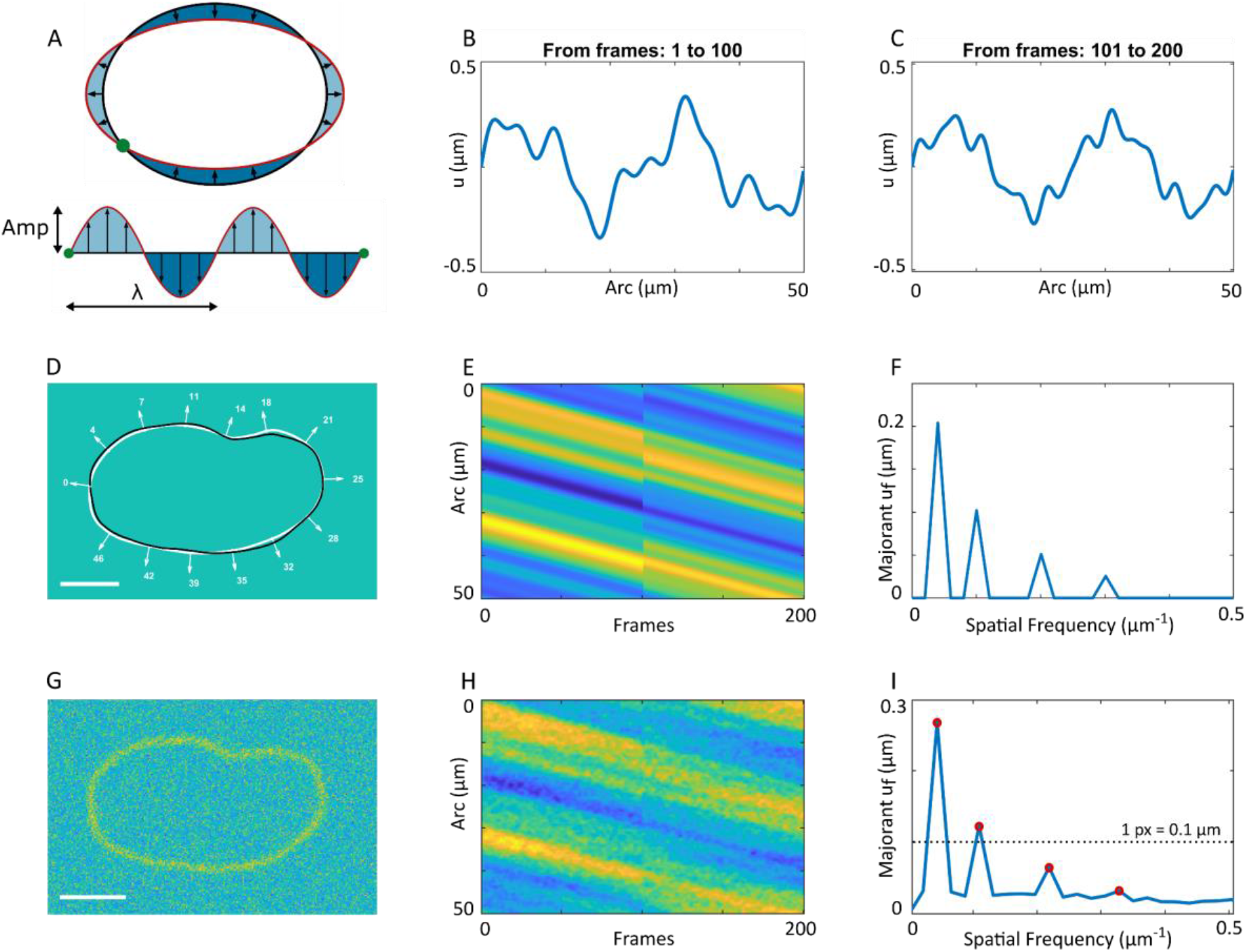
Validation of the membrane fluctuations algorithm using synthetic video. (A) Unfolding the distance between the current membrane (red) and the reference membrane (black) generates a waving signal (bottom) with characteristic wavelengths. For a compression fluctuation, the associated unfolded fluctuations have a wavelength (λ) of half perimeter. (B) Unfolded fluctuation used between frames 1 and 100 to create the synthetic validation video. The full synthetic video has a total of four sinusoidal components: f_1_ = 0.05 μm^−1^, Amp_1_ = 0.2 μm; f_2_ = 0.1 μm^−1^, Amp_2_ = 0.1 μm; f_3_ = 0.2 μm^−1^, Amp_3_ = 0.05 μm; f_4_ = 0.3 μm^−1^, Amp_4_ = 0.025 μm. The fluctuations created between frames 1 and 100 have the components: {f_1_, Amp_1_}, {f_2_, Amp_2_} and {f_3_, Amp_3_}. (C) Unfolded fluctuation used between frames 101 and 200. The signal has three sinusoidal components: {f_1_, Amp_1_}, {f_3_, Amp_3_} and {f_4_, Amp_4_}. (D) First frame of synthetic membrane (back), waving around the median membrane (white). E) Expected fluctuation map of the entire video. The fluctuations (B, C) added to the reference membrane were continuously propagating around the nucleus in order to create a dynamic membrane. This generates the descending pattern shown in E. (F) Majorant of frequency depend fluctuations, *u_f_*, obtained for the map E. The peaks of this curve reveal the amplitude and wavelengths of the sinusoidal waves used to create the fluctuations. (G) Final synthetic video created by adding noise to the original video (D). (H) Fluctuations map obtained by analyzing the noisy video (G) with Trackosome. The map is very similar to the expected result (E), clearly showing the two different fluctuation patterns. (I) Majorant *u_f_* obtained with Trackosome for the noisy video (G), revealing the four expected peaks: f_1_* = 0.044 μm^−1^, Amp_1_* = 0.26 μm; f_2_* = 0.11 μm^−1^, Amp_2_* = 0.12 μm; λ_3_* = 0.22 μm^−1^, Amp_3_* = 0.065 μm; λ_4_* = 0.33 μm^−1^, Amp_4_* = 0.032 μm. Trackosome is able to detect fluctuations with amplitudes at the subpixel level.

**Figure S7.**
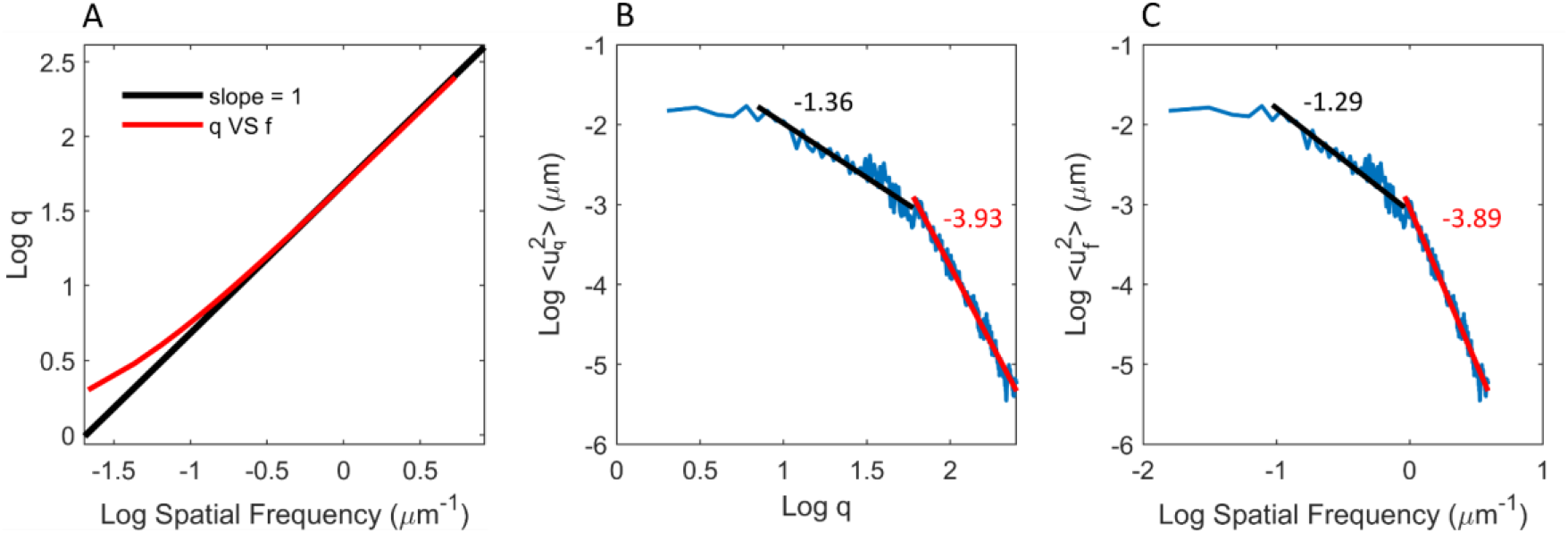
Equivalence between slopes using wavenumber (q) and Spatial frequency. (A) In logarithmic scales, q and Spatial frequency are equivalent for frequencies above 0.1 μm^−1^, making the slope analysis viable for the considered range. (B) Representative example of the average FT of the squared fluctuations, <*u_q_^2^*>, for a given cell plotted against q, and the slopes of the linear fits (solid lines) at low and high frequencies. (C) Same <*u_f_^2^*> curve as in B, but plotted against the associated spatial frequency. The slopes of the linear fits are very similar to those obtained in B.

